# REGULATION OF COLONIC MACROPHAGES AND TYPE-17 AND REGULATORY T CELLS IN DSS-COLITIS BY IBD-ASSOCIATED TRANSCRIPTION FACTOR, CREM

**DOI:** 10.64898/2026.01.26.701728

**Authors:** Shelby L. Schenck, Md Jashim Uddin, Christopher F. Pastore, Audrey C. Brown, William A. Petri

## Abstract

**Background:** Recent genome wide association studies (GWAS) performed by our laboratory identified polymorphisms at the locus containing the gene, cAMP-responsive element modulator (CREM), that influence *Entamoeba histolytica*^+^ diarrheal disease susceptibility in children. CREM is a cAMP-responsive transcription factor that regulates genetic expression and epigenetic modulation in a context- and cell-specific manner. Polymorphisms at this locus have been previously associated with IBD susceptibility, suggesting CREM regulates enteric inflammation in infectious and autoimmune colitis.

**Methods:** Mice were generated with either a tamoxifen-inducible global deletion or an intestinal epithelial cell (IEC)-specific deletion of *Crem*. Dextran-sodium sulfate (DSS) was administered to chemically induce colitis and mice were assayed for weight loss, clinical score, spectral flow cytometry of colonic lamina propria and mesenteric lymph node white blood cells, and shallow shotgun whole genome sequencing of fecal samples.

**Results:** Tamoxifen-inducible global deletion of *Crem* significantly ameliorated DSS-colitis severity as measured by clinical scoring and weight loss over the course of disease (p = 2.29 × 10^-15^, p = 2.24 × 10^-21^, respectively). Protection was not phenocopied when *Crem* was deleted exclusively in IECs. When sampled during acute colitis, protection seen in *Crem*-deleted mice was associated with a significant increase in macrophages, and RORγt^+^ regulatory (pTregs) and T helper (Th17) cells in the colonic lamina propria, along with an increase of T-follicular like helper cells in the mesenteric lymph node.

**Conclusions:** Inducible global deletion of *Crem* reduced the severity of DSS colitis while increasing colonic macrophages, RORγt^+^ regulatory (pTregs) and T helper (Th17) cells. Future work will investigate the aforementioned cell types to determine the mechanism by which *CREM* aggravates DSS-colitis, thereby defining the immunoregulatory role of *CREM* in intestinal inflammation with the goal of identifying new therapeutic targets for IBD.

## INTRODUCTION

2.1 million Americans are afflicted with inflammatory bowel disease (IBD), and its main subtypes, Crohn’s disease (CD) and ulcerative colitis (UC), are among the top ten most prevalent autoimmune diseases.^1^ Inflammation in IBD derives from an uncontrolled and improper immune response to commensal microbiota and/or the tissue itself. Current treatments are variable in efficacy; 90% of patients relapse over a ten-year period despite use of the biologic anti-TNF. Despite advances such as IL-23 inhibition, risankizumab achieved adjusted clinical remission in only 21–22% of patients at 12 weeks in Phase III trials,^2^ underscoring the need for continued research and more effective therapies. Furthermore, complications from insufficiently controlled IBD can be dire, leading to malnutrition^3^ and bowel stricture^4^ or fistula formation^5^ in CD patients, whereas UC complications are commonly bowel perforations or toxic colon enlargement^6^ and increased susceptibility to colon cancer.^7^

In efforts to understand the genetics of the disease, variants of more than 240 gene loci have been associated with IBD susceptibility, 30 of which have been causally linked.^8^ Of these, about 200 have been identified by genome wide association studies (GWAS), where the remainder have been derived through familial studies. The biological relevance of this is exemplified by *Nod2*, a gene that normally recognizes bacterial cell wall components. Single-nucleotide polymorphisms (SNPs) in *Nod2* are found in 30 to 40% of CD patients. ^9^ Recently through GWAS, our laboratory uncovered risk polymorphisms within the *CUL2-CREM-CCNY* locus that influence susceptibility to *Entamoeba histolytica (E. his)*^*+*^ diarrheal disease within the first year of life.^10^ This locus exists in a state of high linkage disequilibrium, preventing identification of a putatively causative SNP. However, additional analysis of eQTLs identified by the Genotype-Tissue Expression (GTEx) Consortium showed that polymorphism(s) implicate *cAMP-responsive element modulator (CREM*) as the likely driver of *E. his*^*+*^ diarrheal disease association. Polymorphisms at the *CREM* locus have also previously been associated with IBD susceptibility including considerable overlap with those identified in *E. his*^*+*^ diarrheal disease GWAS ^11–14^ (rs34779708 p = 2 × 10^-25^; rs12242110, 1 × 10^-9^)^10^. Therefore, *CREM* plausibly regulates enteric inflammation in both autoimmune and infectious colitis.

CREM belongs to the activating transcription factor (ATF) family of transcription factors. CREM has several isoforms due to alternative splice sites and multiple promoters, leading to both activating and suppressing isoforms ^15^. The *CREM* transcription factor is broadly expressed across human tissues^16^ and immune cells^17–20^ Functionally, CREM binds to the cAMP response element (CRE) sequence following cAMP-dependent phosphorylation or calcium signaling. CREM additionally regulates transcription by recruiting other transcription factors and/or by the recruitment of epigenetic modulators such as histone deacetylase (HDAC) and DNA methyltransferase (DMNT).^21–23^ Notable CREM isoforms include inducible cAMP early repressors (ICERs) and CREMα.

Though the original body of CREM literature focuses on its role in spermatogenesis, recent work has focused on the impact of CREM in immunoregulation. Both ICER and CREMα isoforms have been shown to limit T cell activation. Specifically, CREM acts in a negative feedback look following AP-1/cFos activation following *in vitro* T cell stimulation.^24^ Additionally, following phosphorylation by CaMK4, CREM can further suppress T cell activation by recruiting HDAC1 and DMNT3a to the IL2 promoter, suppressing IL2 expression.^25^ This negative-feedback loop exists in other innate immune cells as well, where CREM increases following IL-15R downstream signaling in NK cells.^26^ Similarly, high levels of CREM has also been associated with an immunosuppressive and highly macrophage-infiltrated microenvironment in gastric cancer,^27^ supporting a role for CREM in dampening immunity.

Despite the characterized immunosuppressive capacity of CREM, ICER protein levels and *CREMα* mRNA levels are significantly increased in the T helper (Th) cells of patients with systemic lupus erythematosus (SLE) compared to healthy controls, ^25,28,29^ suggesting a role for CREM in promoting autoimmunity. This has been echoed in several autoimmune mouse models, including glomerulonephritis and SLE, where loss of CREM is protective^28^. Of relevance in mucosal immunology, *CREM* expression is higher in patients with UC and is expressed at high levels in CD4 and CD8 T cells and monocytes.^30^

We hypothesized that *CREM* regulates colonic inflammation in autoimmune colitis. To understand how CREM affects IBD pathology, we investigated the effect of inducible global deletion of *Crem* during DSS-colitis. To identify how *Crem* expression alters the immunobiology of DSS-colitis, we assessed disease progression by weight loss and qualitative clinical scoring, and we conducted spectral flow cytometry of colonic lamina propria and mesenteric lymph node (MLN) populations. Here, we show that global loss of *Crem* protects mice from DSS-colitis weight loss and disease severity score. Additionally, protected *Crem* global deletion mice have significantly higher intestinal macrophages and type-17 and regulatory T cells. Additionally, these protected mice have significantly higher levels of T follicular-like helper cells within the mesenteric lymph node. Together, our work suggests that *Crem* promotes DSS-colitis and identifies potential cellular arbiters, thereby offering insight into the immunobiology underlying the genetic association between *CREM* and enteric immunity.

## RESULTS

### Inducible global deletion of Crem ameliorates DSS-colitis severity

To test if the expression of *Crem* affects DSS-colitis, *Crem*^*fl/fl*^ mice with LoxP sites flanking the basic domain of *Crem* were produced an a C57Bl/6 background^31^ **(Figure 1A)**. Expression of Cre recombinase would therefore delete the basic domain of CREM, which recognizes the cAMP response element (CRE) of DNA (5’ – TGACGTCA – 3’). The basic domain is conserved between humans and mice and is required for function of all known CREM isoforms.^3^ *Crem*^fl/fl^ mice were bred to *Cre-ER*^32^ mice to generate inducible global deletion model of *Crem*, which we confirmed previously.^31^ Mice were administered tamoxifen to induce global deletion of *Crem* prior to treatment with 2.5% dextran sodium sulfate (DSS) **(Figure 1B)**. Mice deficient in *Crem* lost significantly less weight (*p = 2.24* × *10^-21^*) and were significantly lower in qualitative clinical scoring (*p = 2.29* × *10^-15^*) compared to same-sex floxed littermate controls throughout the experiment **(Figure 1C-D)**. However, when mice were euthanized on day 7, where there was the greatest separation between groups, there was no significant difference in colon length. This suggests that *Crem* promotes DSS-colitis, potentially via the gut response to DSS-induced damage and inflammation rather than lessening the impact on the epithelium or fibrosis. Additionally, it is possible that there may be differences in colon length during recovery rather than our current sampled acute timepoint.

**Figure 1.**
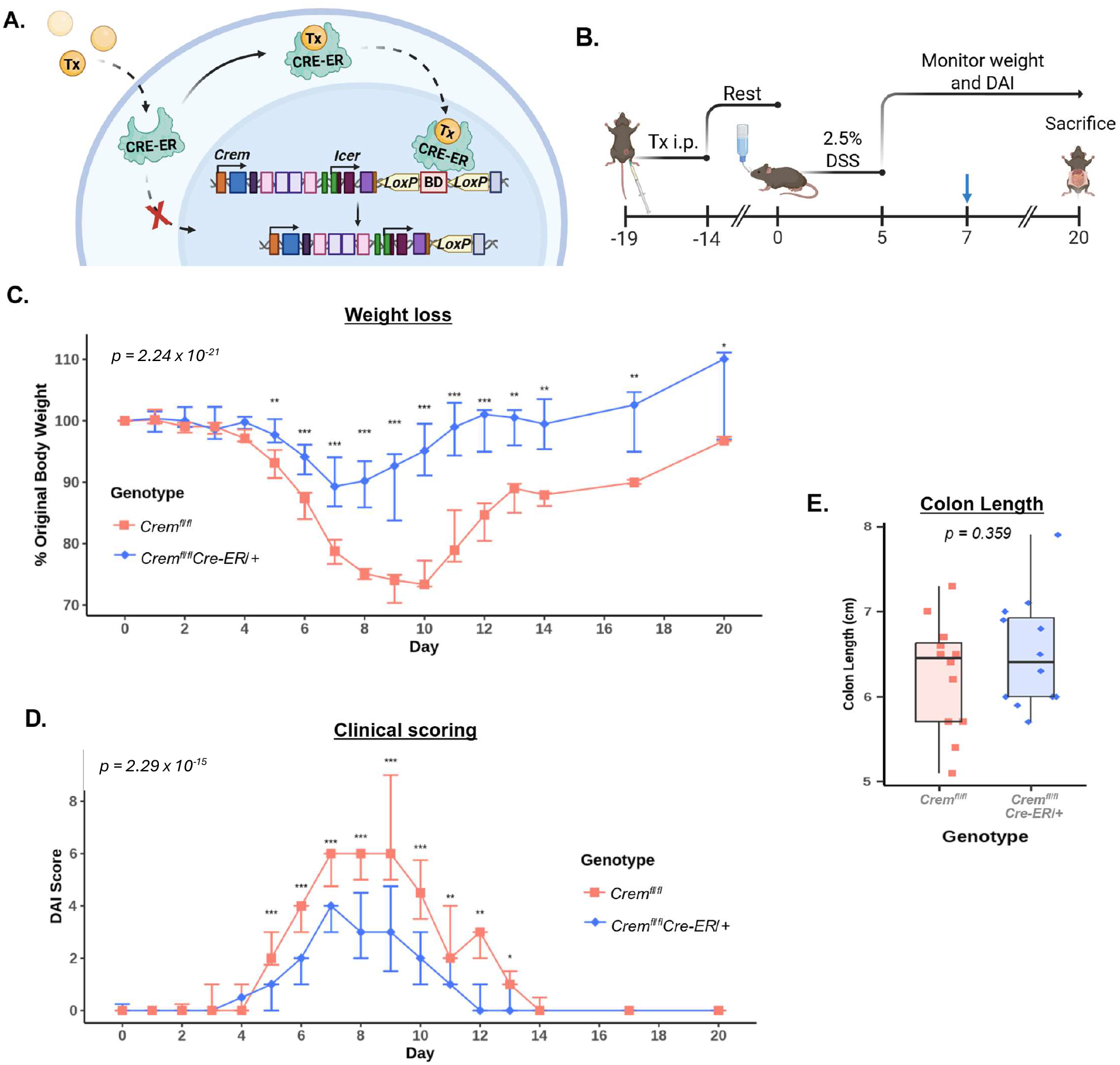
Inducible global Crem deletion protects against acute DSS-colitis. **A**. Schematic of tamoxifen (Tx) inducible deletion system. **B**. Schematic of experimental timeline. Blue arrow indicates day mice were harvested to obtain colons for *E* and flow cytometry experiments. Tx, Tamoxifen; DAI, disease activity index. **C**. Percent weight loss of *Crem*^*fl/fl*^ and *Crem*^*fl/fl*^*Cre-ER/+* mice during DSS-colitis. **D**. Disease activity index of mice during DSS-colitis, scored on diarrhea, posture, and activity. **C-D**. Combined data from three independent experiments are presented as median + IQR, n = 16 per group. Statistics are calculated from a Linear Mixed Model, where % Original Body Weight or DAI is explained by an interaction between fixed effects, genotype and day, and random effect being the observation per mouse. Model p values were generated by Type III ANOVA. **E**. Length of colon at 7d of DSS-colitis. Combined data from two independent experiments are presented as median ± IQR, n = 11 per group. * p ≤ 0.05, ** p ≤ 0.01, and *** p ≤ 0.001.

### *Crem* expression in intestinal epithelial cells does not impact DSS-colitis

*Crem* is expressed throughout the gastrointestinal tract.^16^ DSS induces inflammation by damaging the single layer of epithelium separating the lamina propria from the gut lumen, inducing a state of dysbiosis and allowing translocation of potentially inflammatory microbes and food antigens. *F*ollowing antibiotic-induced gut dysbiosis, *Crem*-regulated transcriptional network, or regulon, increases in intestinal epithelial cells (IECs.)^33^ This suggests *Crem* may protect from DSS-colitis by modulating the IEC response to DSS-induced dysbiosis. Moreover, ICER has been shown to promote apoptosis by inhibition of Bcl6, an anti-apoptotic protein,^34,35^ suggesting that the loss of *Crem* within IECs may protect from DSS by maintaining the gut barrier through lessened damage-induced apoptosis.. To test if *Crem* in IECs promotes DSS-colitis, we crossed our *Crem*^*fl/fl*^ mice to *Vil*^*Cre* 36^to generate conditional IEC-specific deletion of *Crem* and confirmed as described.^31^ Mice were challenged with 2.5% DSS; however, IEC-specific deletion of *Crem* did not phenocopy protection seen in inducible global deletion **(Supp Fig 1A-B)**. Therefore, it is unlikely that *Crem* acts within IECs to promote DSS-colitis.

### Immunophenotyping of inducible global Crem deletion in DSS-colitis

Functionally-relevant *CREM* expression has been demonstrated in several immune cells^17–20^. scRNAseq has demonstrated that *CREM* is higher in the gut of patients with UC compared to healthy controls, particularly in CD4 and CD8 T cells and monocytes.^30^ Additionally, *Crem* has been shown to poise T cells towards IL-17 production^28,37^ and metabolically inhibits the regulatory T cell phenotype ^38^. The Th17/Treg ratio has often been interrogated in the context of several IBD mouse models, including DSS.^39,40^ Therefore, we asked if loss of *CREM* influences immune cell populations during peak DSS-colitis within the colonic lamina propria and the mesenteric lymph node (MLN), with an interest in type-17 and regulatory T cells. Day 7 was chosen as the optimal timepoint to capture differences between the control and inducible global *Crem* deletion groups as this timepoint had the greatest disparity in weight loss and disease severity **(Fig 1C-D)**.

### Inducible global deletion of Crem increases type-17 and regulatory T cells and macrophages in the colonic lamina propria

There was a significant increase in both regulatory and type-17 T cells. Specifically, the percent of CD4^+^ T cells that were FOXP3^+^ (regulatory T cells, Tregs), FOXP3^+^ RORγt^+^ (peripheral Tregs, pTregs), and FOXP3^-^ RORγt^+^ (T helper 17, Th17) was increased in the colon lamina propria of inducible global *Crem* deletion mice **(Figure 4A)**. The significant increase in both pTregs and Th17 cells was maintained by percentage of live cells as a measure of count **(Figure 4B)**. Additionally, macrophages quantified as the percentage of CD45^+^ or of live cells were significantly increased in these deletion mice **(Figure 4C-D)**. Population increases demonstrated by manual gating were largely echoed by Uniform Manifold Approximation and Projection (UMAP), where density plots of CD45^+^ colonic lamina propria cells from inducible global *Crem* deletion mice showed a visible increase in regulatory T cells and macrophages **(Figure 2)**. Though statistical analysis of our current panel did not reveal any significant differences in dendritic cells (DC), FlowSOM cluster analysis demonstrated a trending increase in DC subsets, clusters 6 and 8, in the inducible global *Crem* deletion mice **(Figure 2B)**. Therefore, *Crem* may promote DSS-colitis through macrophages and/or type-17 and regulatory T cells.

**Figure 2.**
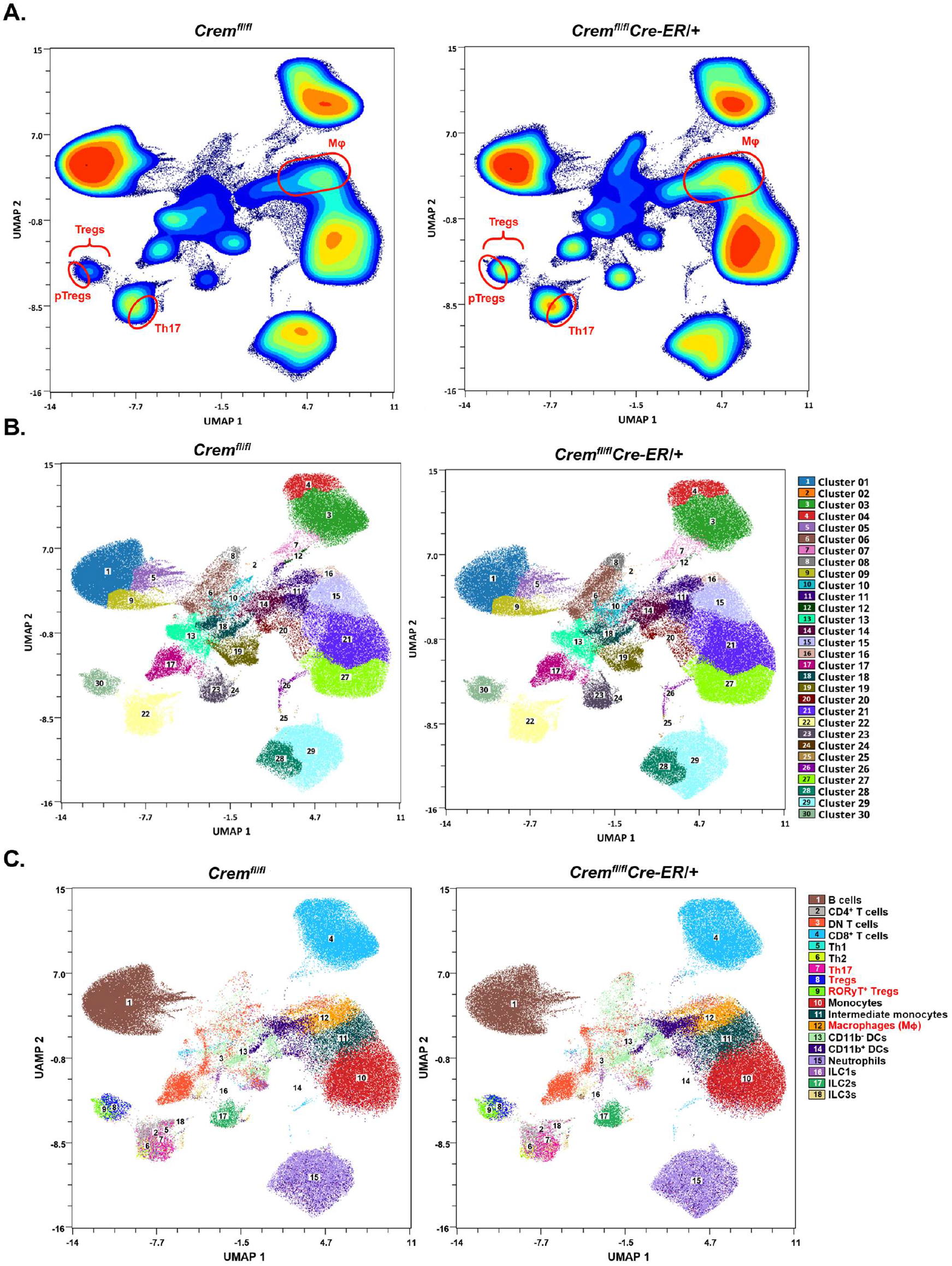
UMAP of colonic immune populations by spectral flow cytometry shows enrichment of regulatory T cells and macrophages in inducible global Crem deletion mice. UMAP of three-representative mouse colons displayed as **A**. density plot **B**. cluster overlay, and **C**. manual gating. **(A, C.)** Populations of interest are colored in red and highlighted where applicable.

### Inducible global Crem deletion increases T follicular helper cells in the mesenteric lymph node

Within the MLN, the percentage CD4^+^ T cells that are CXCR5^+^ (T follicular helper cells, Tfh) was also significantly increased **(Figure 4E)**. Furthermore, this phenotype trended strongly by count as quantified by percentage of live cells (**Figure 4F)**. Additional analysis by UMAP did not reveal any shifts in additional populations and recapitulated the increase seen in Tfh cells by density plot **(Figure 3)**. These data therefore suggest that *Crem* may have an impact on humoral and cellular immunity in the context of DSS-colitis.

**Figure 3.**
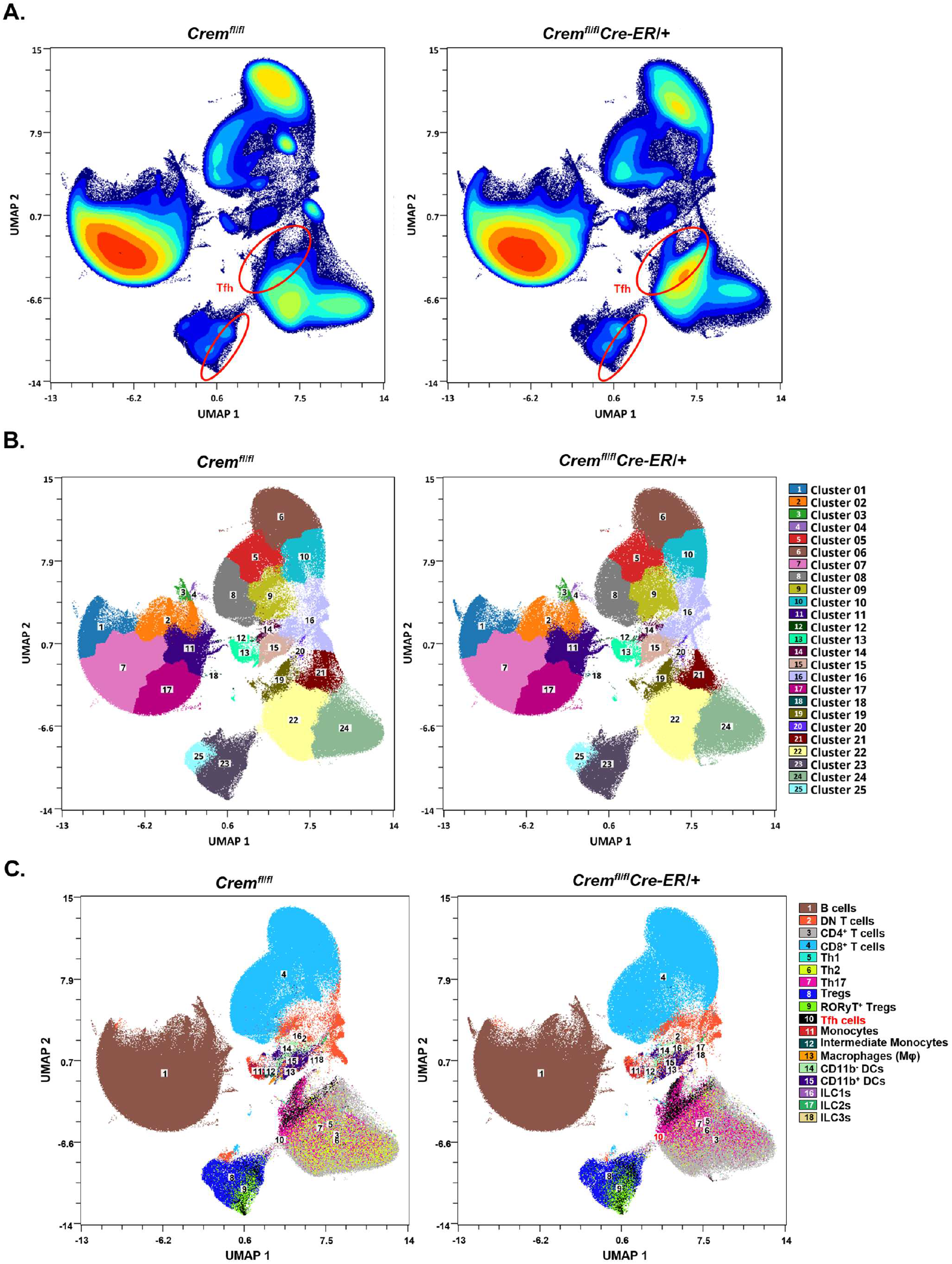
UMAP of mesenteric lymph node immune populations by spectral flow cytometry shows enrichment of T follicular helper T cells in inducible global Crem deletion mice. UMAP of three-representative mouse mesenteric lymph nodes (MLN) displayed as **A**. density plot **B**. cluster overlay, and **C**. manual gating. **(A, C.)** Populations of interest are colored in red and highlighted where applicable.

**Figure 4.**
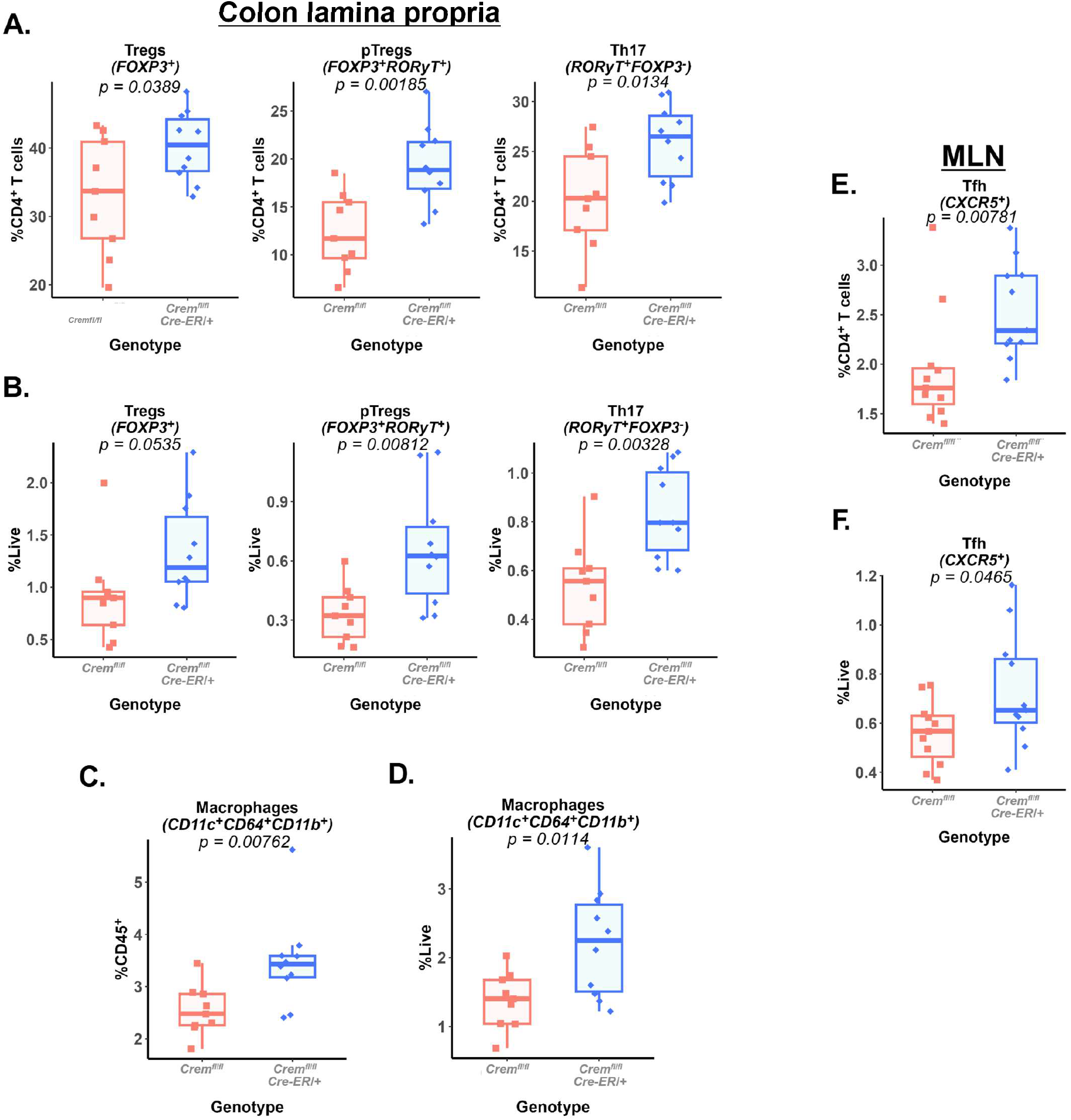
Inducible global Crem increases regulatory and type-17 T cells and macrophages in the colon, and T follicular helper cells in the MLN. **A-B**. Percent of colon lamina propria *(A)* CD4^+^ T cells *(B)* live cells that are regulatory T cells (Tregs, FOXP3^+^, left), peripheral T regulatory T cells ^or^ (pTregs, FOXP3^+^ RORγT^+^, middle), T helper 17 (Th17, RORγT^+^ FOXP3^-^, right). **C-D**. Percent of colon lamina propria *(C)* CD45^+^ or *(D)* live that are macrophages (CD11c^+^ CD64^+^ CD11b^+^). **E-F**. Percent of mesenteric lymph node (MLN) *(E)* CD4^+^ T cells or *(F)* that are T follicular helper cells (Tfh, CXCR5^+^). **A-F**. *(A-D)* n = 9 *Crem* ^*fl/fl*^ and n = 10 *Crem*^*fl/fl*^*Cre-ER/+* and *(E-F)* n = 11 per group. Statistics were calculated as *(A, B - middle, right, F)* Student’s T test, *(B - left, C, E,)* Mann Whitney U Test, and *(D)* unequal T test. Data presented as median ± IQR.

### Crem does not impact the fecal bacteriome at steady-state or during DSS-induced dysbiosis

pTregs and Th17 cells are dependent on the microbiota both in the context of differentiation and function^41–44^. Additionally, DSS-colitis is microbiome dependent; for example, germ-free mice and antibiotic-treated mice possess significantly less inflammation under DSS treatment compared to conventional specific pathogen free mice.^45,46^ Therefore, to determine if *Crem* impacts the microbiome during DSS-induced dysbiosis, we conducted shallow whole-genome shotgun sequencing at baseline (Day 0) and peak disease severity (Day 7) of DSS colitis **(Supp Fig 2A)**. Alpha diversity decreased after DSS treatment as seen previously^47^; however, there was no difference between mice lacking *Crem* and their same-sex littermates before or during DSS-colitis **(Supp Fig 2B)**. Additionally, there was no difference in relative taxa abundance **(Supp Fig 2C)** during DSS colitis. Beta diversity grouped samples by timepoint irrespective of *Crem*, and there was little to no differences in top contributors to PC2 to parse out the minimal separation of inducible global *Crem* deletion mice and same-sex littermates **(Supp Fig 2D)**. Therefore, *Crem* did not impact the bacteriome at steady-state or during DSS-colitis.

## DISCUSSION

We demonstrate that *Crem* acts not only as a biomarker for IBD susceptibility but regulates IBD immunobiology. Moreover, loss of *Crem* significantly protected mice from DSS-colitis severity and was associated with a significant increase in type 17 and regulatory T cells and macrophages in the colonic lamina propria. Additionally, this phenotype coincided with a significant increase in Tfh cells in the MLN. This highlights a potential mechanistic role for *Crem* in the context of IBD, echoing previous work that supports the a deleterious role for increased CREM expression in autoimmunity in people and several mouse models.^25,28,48^

### Crem may inhibit anti-inflammatory macrophages

Macrophages are one of the earliest effectors in DSS-colitis, responding indiscriminately to self- and nonself-antigens within the colon lamina propria. *Crem* expression increases downstream of toll-like receptor (TLR) activation^20^ and is differentially expressed in macrophages from germ-free and conventional mice,^19^ suggesting *Crem* may regulate the macrophage response to the microbiome. We showed that inducible global deletion of *Crem* protects mice from DSS-colitis and is associated with an increase in macrophages. This is supported by work where increased CREM expression is associated with macrophages in patients with cystic fibrosis^49^ and monocytes in patients with UC.^30^ Anti-inflammatory M2 IL10R^+^ macrophages are essential for IL10-mediated protection of DSS-colitis.^50^ These IL10R^+^ macrophages have also been demonstrated to be essential in establishment of anti-inflammatory Th17 cell niche.^42^ Additionally, IL10-sensing CD206^+^ macrophages have been suggested to work in a positive-feedback loop to promote Tregs post recruitment via the CCL7/8:CCR2/5 axis.^51^ As CREM has been shown to be differentially expressed in macrophages of germ-free versus conventional mice,^19^ it is plausible that CREM may impact both the macrophages response to the microbiome and subsequently the anti-inflammatory Th17/Treg niche.

### Crem may inhibit T cell-mediated immunosuppression in DSS-colitis

IBD is characterized by loss of or improper tolerogenic response in the gut. Previous work has shown that T cell-specific overexpression of the *CREMα* isoform is deleterious in chemically-induced colitis and associated with a local increase in IL-21^52^. Concordantly, we showed that the protective *Crem*-deficient DSS-colitis was associated with an increase in total Tregs and pTregs. Unlike thymic-derived or naturally-derived Tregs (nTregs), pTregs are tolerized within peripheral tissues. These cells are thought to represent a more stable regulatory T cell subset compared to nTregs and are characterized by increased demethylation at *Foxp3 and Ctla4*. ^53^ Notably, pTregs associated with a greater suppressive capacity than nTregs^53^, suggesting they may more strongly promote tolerance. Furthermore, pTregs, which express RORγT^+^, are partly type-17 polarized and are associated with tissue regeneration^54^. Therefore, pTregs may both dampen acute disease and strength repair in *Crem*-deficient DSS-colitis.

Interestingly, in *Crem*-deficient DSS-colitis, the increase in regulatory T cells coincided with an increase in Th17 cells. This is contrary to the established literature on IBD and DSS-colitis where Tregs and Th17s often act in opposition to each other.^39,55–58^ However, IL-17 blockade has been unsuccessful in effectively treating IBD and at times exacerbates disease.^59,60^ This may be because canonical type-17 cytokines like IL22 and IL17A are capable of preserving barrier integrity and limiting bacterial dissemination.^61–63^ Additionally, Th17 cells can be induced to an anti-inflammatory subtype in a microbiota-dependent manner and can subsequently exhibit a strong suppressive capacity.^42^ Therefore, this positive association of Th17 cells with protection during *Crem*-deficient DSS-colitis phenotype plausibly suggests that they may act alongside Tregs and pTregs to mitigate inflammation.

### Crem may oppose humoral immunity in DSS-colitis

Herein, we demonstrate that protective *Crem* deletion during DSS-colitis is associated with a significant increase in Tfh cells within the MLN. Common antibodies in IBD include those against intestinal microbes, like anti-*S. cerevisiae* and anti-flagellin, and those against self, such as perinuclear anti-neutrophil cytoplasmic antibodies (pANCA). The former are found more commonly in CD patients (∼65% and 55% of cases, respectively), while the later are found in a 65% of UC patients.^64^ Though presence of these antibodies are not a crucial diagnostic criterion, they may be predictive of response to treatment as patients that are pANCA-positive show lower efficacy in response to TNF-inhibition, one of the leading biologics to treat IBD.^65–67^ High-affinity pANCAs have been correlated with worsened disease in ANCA-associated vasculitis;^68,69^ similarly, high affinity and avidity antibodies have been shown to mark colitogenic bacteria.^70^ Therefore, as our data demonstrated an increased Tfh cells in protected mice following *Crem* deletion, it is possible *Crem* may antagonize production of low-affinity antibodies, thereby lessening immune exclusion of microbial- and self-antigens released during DSS-induced injury.

A limitation of this study is that, while we demonstrate that inducible global deletion of *Crem* protects mice from weight loss and disease severity in the well-characterized IBD animal model, DSS-colitis, this does not recapitulate previous eQTL data from GTEx predicted that IBD risk-associated *CREM* alleles lower *CREM* expression.^10^ Moreover, previous work from our laboratory showed small intestinal biopsies stratified by genotype for a SNP at this locus did not find differential expression of *CREM*.^31^ Although allele-specific editing would create an ideal model, extensive linkage disequilibrium at the CREM locus prevents identification of putatively causal genetic variant(s). Future work to conduct a fine-mapping analysis of the CUL2-CREM-CCNY locus may aid the identification of a causal SNP for IBD susceptibility. However, our data that loss of *Crem* was protective in DSS-colitis is supported by previous work where UC patients were shown to have significantly higher expression of *CREM* in the gut compared to healthy controls.^30^

Our work revealed that inducible global deletion of *Crem* protects mice from DSS-colitis and increases the percentage of several known players in IBD and DSS-colitis immunopathology. Future work will test if *Crem* inhibits anti-inflammatory macrophages, thereby lessening the establishment of tolerogenic pTregs/Th17 niche. Additional work will examine if *Crem* inhibits Tfh cells, thus limiting B cell production of low-affinity secretory IgA and subsequent commensal immune exclusion.

## MATERIALS AND METHODS

### Mice

All mouse experiments adhered to the ethical guidelines for animal research and conducted under protocols approved by the Institutional Animal Care and Use Committee (IACUC) at the University of Virginia. Experiments utilized same-sex littermate when able or sex/age-matched, aged between 6 to 25 weeks at start of DSS. Mouse genotypes from tail biopsies were determined using real-time PCR with probes specifically designed for each line (Transnetyx, Cordova, TN). All animals were housed in a specific pathogen-free environment at the animal facility at the University of Virginia.

### Inducible global deletion of Crem

*Crem*^*fl/fl*^, *Crem*^*fl/fl*^*Cre-ER/+*, and *Crem*^*fl/fl*^*Vil1*^*Cre/+*^ mice were originally generated and confirmed as previously described.^31^ *Crem*^*fl/fl*^*Cre-ER/+* mice and their respective same-sex littermate or sex/age-matched floxed control mice, *Crem*^*fl/fl*^, were given tamoxifen (Sigma-Aldrich T5648) for 5 d (20mg/mL in corn oil [Sigma-Aldrich C8267] 100 μL) intraperitoneally and allowed to rest for approximately two weeks to induce Cre activity and subsequent *Crem* deletion, as determined previously.^31^

### Experimental colitis

Mice were administered 2.5% dextran sodium sulfate (DSS; 40-50 kD, Thermo Scientific J14489) in their drinking water for 5 days to induce acute colitis. Afterwards, mice were transitioned to regular water. Mice were weighed and scored daily during this period and up to day 14. Then mice were weighed and scored every third day until day 20. Mice were evaluated by clinical scoring of the following parameters: weight loss, activity level, stool consistency, and posture. Each parameter was scored and added to a cumulative clinical score ranging 0 to 20. Weight loss and activity scores ranged 0 to 4; Stool consistency and posture were scored from 0 to 3 **(Supp Table 2)**. Mice with a weight loss score of 4 (greater than 25%), an activity score of 4, or a clinical score of 15 or higher were euthanized as per animal protocol. All experiments were conducted with DSS-treated mice unless otherwise specified.

### Spectral flow cytometry

Mice were humanely euthanized by carbon dioxide asphyxiation followed by cervical dislocation on day 7. Colon and mesenteric lymph node (MLN) were resected and placed into buffer (5% FBS and 25mM HEPES in HBSS) or RPMI complete (10% FBS in RPMI 1640), respectively, and kept on ice. Colon epithelium was separated from the lamina propria by incubating for 40 minutes in dissociation buffer (15 mM HEPES, 5 mM EDTA, 10% FBS, and 1 mM DTT in HBSS) at 37C with gentle shaking. Tissue was then cut into smaller pieces and digested using digestion buffer (0.17 mg/mL Liberase TL and 30 μg/mL DNase in RPMI 1640). Cells were then passed sequentially through 100 μM and 40 μM cell strainers to obtain single-cell suspensions in FACS buffer (2% FBS in PBS). Cells were washed with PBS and incubated with Fc block (BioLegend 156604) for 10 minutes at room temperature, washed with PBS, followed by LIVE/DEAD BLUE. Cells were then washed with FACS buffer and stained with extracellular staining for 30 minutes at 4C. Cells were then washed with FACS buffer and permeabilized with Foxp3 Fix/Perm Working Solution (ThermoFisher Scientific 00-5523-00) and incubated at room temperature for 30 minutes. Cells were washed with permeabilization buffer, followed by intracellular staining. Cells were washed with FACS buffer and spectral flow cytometry was performed on the Cytek Aurora (5-laser) Spectral Flow Cytometer and analyzed in OMIQ. Gating strategy and markers are in in the supplement **(Supp Fig 3-4, Supp Table 1**). Statistics were generated using R (version: 4.3.3). Spectral flow cytometry data were subset to include equal numbers of live CD45^+^ single cells from each sample in OMIQ and clustered by FlowSOM for Uniform Manifold Approximation and Projection (UMAP) analysis.

### Microbiome analysis

Stool was collected from mice undergoing experimental colitis every other day from 0d to 7d. Mice were co-housed prior to and throughout sample collection. Stool samples were transferred to TransnetYX microbiome buffer solution upon collection and shipped to TransnetYX for using shallow shotgun whole genome sequencing with a read depth of 2 million paired-end reads. Results were then analyzed through the OneCodex platform. Measures of alpha and beta diversity were downloaded from OneCodex, along with relative taxa abundance. Relative species abundance values were transformed and analyzed using R (version: 4.3.3) by PERMANOVA using the package, vegan (version: 2.6-10^71^), to determine if there was an association with *Crem* expression, state of DSS-colitis, or an interaction between *Crem* and the state of DSS-colitis at collection.

### Statistical analysis

Comparisons involving repeated measures of the two groups, such as weight loss and disease activity index, were generated using linear mixed modeling. Statistical tests for spectral flow cytometry are as described in figure legends. All statistical analyses were performed using R (version: 4.3.3).

## Acknowledgments

We acknowledge members of the Petri laboratory at UVA for technical help and scientific discourse. Additionally, we would also like to acknowledge the UVA Flow Cytometry Core Facility for technical support in this work. BioRender was used to generate schematics in figures. This work was supported by grants from the NIH 5R37AI026649-35 to WAP and a PhRMA Foundation Postdoctoral Fellowship in Translational Medicine to ACB.

## Data Availability Statement

All R code used to generate statistics and analyze data will be uploaded to GitHub, and microbial data will be deposited to an online repository, upon full publication.

## Notes

### Competing Interest Statement

The authors have declared no competing interest.

## REFERENCES

1 Abend AH, He I, Bahroos N, Christianakis S, Crew AB, Wise LM, et al. Estimation of prevalence of autoimmune diseases in the United States using electronic health record data. J Clin Invest n.d.;135:e178722. 10.1172/JCI178722.

2 D’Haens G, Panaccione R, Baert F, Bossuyt P, Colombel J-F, Danese S, et al. Risankizumab as induction therapy for Crohn’s disease: results from the phase 3 ADVANCE and MOTIVATE induction trials. The Lancet 2022;399:2015–30. 10.1016/S0140-6736(22)00467-6.

3 Balestrieri P, Ribolsi M, Guarino MPL, Emerenziani S, Altomare A, Cicala M. Nutritional Aspects in Inflammatory Bowel Diseases. Nutrients 2020;12:372. 10.3390/nu12020372.

4 Bettenworth D, Bokemeyer A, Baker M, Mao R, Parker CE, Nguyen T, et al. Assessment of Crohn’s disease-associated small bowel strictures and fibrosis on cross-sectional imaging: a systematic review. Gut 2019;68:1115–26. 10.1136/gutjnl-2018-318081.

5 Basiji K, Kazemifard N, Farmani M, Jahankhani K, Ghavami SB, Fallahnia A, et al. Fistula in Crohn’s disease: classification, pathogenesis, and treatment options. Tissue Barriers 2025;13:2458784. 10.1080/21688370.2025.2458784.

6 Silverberg D, Rogers AG. TOXIC MEGACOLON IN ULCERATIVE COLITIS. Can Med Assoc J 1964;90:357–63.

7 Zhang L, Zhang X, Su T, Xiao T, Xu H, Zhao S. Colorectal cancer risk in ulcerative colitis: an updated population-based systematic review and meta-analysis. eClinicalMedicine 2025;84:. 10.1016/j.eclinm.2025.103269.

8 El Hadad J, Schreiner P, Vavricka SR, Greuter T. The Genetics of Inflammatory Bowel Disease. Mol Diagn Ther 2024;28:27–35. 10.1007/s40291-023-00678-7.

9 Kayali S, Fantasia S, Gaiani F, Cavallaro LG, de’Angelis GL, Laghi L. NOD2 and Crohn’s Disease Clinical Practice: From Epidemiology to Diagnosis and Therapy, Rewired. Inflammatory Bowel Diseases 2024:izae075. 10.1093/ibd/izae075.

10 Wojcik GL, Marie C, Abhyankar MM, Yoshida N, Watanabe K, Mentzer AJ, et al. Genome-Wide Association Study Reveals Genetic Link between Diarrhea-Associated Entamoeba histolytica Infection and Inflammatory Bowel Disease. mBio 2018;9:e01668–18. 10.1128/mBio.01668-18.

11 Jostins L, Ripke S, Weersma RK, Duerr RH, McGovern DP, Hui KY, et al. Host-microbe interactions have shaped the genetic architecture of inflammatory bowel disease. Nature 2012;491:119–24. 10.1038/nature11582.

12 Anderson CA, Boucher G, Lees CW, Franke A, D’Amato M, Taylor KD, et al. Meta-analysis identifies 29 additional ulcerative colitis risk loci, increasing the number of confirmed associations to 47. Nat Genet 2011;43:246–52. 10.1038/ng.764.

13 Franke A, McGovern DPB, Barrett JC, Wang K, Radford-Smith GL, Ahmad T, et al. Genome-wide meta-analysis increases to 71 the number of confirmed Crohn’s disease susceptibility loci. Nat Genet 2010;42:1118–25. 10.1038/ng.717.

14 Barrett JC, Hansoul S, Nicolae DL, Cho JH, Duerr RH, Rioux JD, et al. Genome-wide association defines more than 30 distinct susceptibility loci for Crohn’s disease. Nat Genet 2008;40:955–62. 10.1038/ng.175.

15 Foulkes NS, Sassone-Corsi P. More is better: Activators and repressors from the same gene. Cell 1992;68:411–4. 10.1016/0092-8674(92)90178-F.

16 Kaprio H, Heuser VD, Orte K, Tukiainen M, Leivo I, Gardberg M. Expression of Transcription Factor CREM in Human Tissues. J Histochem Cytochem 2021;69:495–509. 10.1369/00221554211032008.

17 Sawka-Verhelle D, Escoubet-Lozach L, Fong AL, Hester KD, Herzig S, Lebrun P, et al. PE-1/METS, an antiproliferative Ets repressor factor, is induced by CREB-1/CREM-1 during macrophage differentiation. J Biol Chem 2004;279:17772–84. 10.1074/jbc.M311991200.

18 Lippe R, Ohl K, Varga G, Rauen T, Crispin JC, Juang Y-T, et al. CREMα overexpression decreases IL-2 production, induces a T(H)17 phenotype and accelerates autoimmunity. J Mol Cell Biol 2012;4:121–3. 10.1093/jmcb/mjs004.

19 Kang B, Alvarado LJ, Kim T, Lehmann ML, Cho H, He J, et al. Commensal microbiota drive the functional diversification of colon macrophages. Mucosal Immunol 2020;13:216–29. 10.1038/s41385-019-0228-3.

20 Harzenetter MD, Novotny AR, Gais P, Molina CA, Altmayr F, Holzmann B. Negative Regulation of TLR Responses by the Neuropeptide CGRP Is Mediated by the Transcriptional Repressor ICER1. The Journal of Immunology 2007;179:607–15. 10.4049/jimmunol.179.1.607.

21 Tenbrock K, Juang Y-T, Leukert N, Roth J, Tsokos GC. The transcriptional repressor cAMP response element modulator alpha interacts with histone deacetylase 1 to repress promoter activity. J Immunol 2006;177:6159–64. 10.4049/jimmunol.177.9.6159.

22 Rauen T, Hedrich CM, Juang Y-T, Tenbrock K, Tsokos GC. cAMP-responsive element modulator (CREM)α protein induces interleukin 17A expression and mediates epigenetic alterations at the interleukin-17A gene locus in patients with systemic lupus erythematosus. J Biol Chem 2011;286:43437– 46. 10.1074/jbc.M111.299313.

23 Hedrich CM, Crispín JC, Rauen T, Ioannidis C, Koga T, Rodriguez Rodriguez N, et al. cAMP Responsive Element Modulator (CREM) α Mediates Chromatin Remodeling of CD8 during the Generation of CD3+CD4−CD8− T Cells. J Biol Chem 2014;289:2361–70. 10.1074/jbc.M113.523605.

24 Rauen T, Benedyk K, Juang Y-T, Kerkhoff C, Kyttaris VC, Roth J, et al. A novel intronic cAMP response element modulator (CREM) promoter is regulated by activator protein-1 (AP-1) and accounts for altered activation-induced CREM expression in T cells from patients with systemic lupus erythematosus. J Biol Chem 2011;286:32366–72. 10.1074/jbc.M111.245811.

25 Juang Y-T, Wang Y, Solomou EE, Li Y, Mawrin C, Tenbrock K, et al. Systemic lupus erythematosus serum IgG increases CREM binding to the IL-2 promoter and suppresses IL-2 production through CaMKIV. J Clin Invest 2005;115:996–1005. 10.1172/JCI22854.

26 Rafei H, Basar R, Acharya S, Hsu Y-S, Liu P, Zhang D, et al. CREM is a regulatory checkpoint of CAR and IL-15 signalling in NK cells. Nature 2025;643:1076–86. 10.1038/s41586-025-09087-8.

27 Yu K, Kuang L, Fu T, Zhang C, Zhou Y, Zhu C, et al. CREM Is Correlated With Immune-Suppressive Microenvironment and Predicts Poor Prognosis in Gastric Adenocarcinoma. Front Cell Dev Biol 2021;9:697748. 10.3389/fcell.2021.697748.

28 Yoshida N, Comte D, Mizui M, Otomo K, Rosetti F, Mayadas TN, et al. ICER is requisite for Th17 differentiation. Nat Commun 2016;7:12993. 10.1038/ncomms12993.

29 Lippe R, Ohl K, Varga G, Rauen T, Crispin JC, Juang Y-T, et al. CREMα overexpression decreases IL-2 production, induces a T(H)17 phenotype and accelerates autoimmunity. J Mol Cell Biol 2012;4:121–3. 10.1093/jmcb/mjs004.

30 He Z, Zhou Q, Du J, Huang Y, Wu B, Xu Z, et al. Integrated single-cell and bulk RNA sequencing reveals CREM is involved in the pathogenesis of ulcerative colitis. Heliyon 2024;10:e27805. 10.1016/j.heliyon.2024.e27805.

31 Brown AC, Uddin MdJ, Munday RM, Naz F, Moreau GB, Ramakrishnan G, et al. The cAMP responsive element modulator (CREM) transcription factor influences susceptibility to undernutrition and infection. mBio 2025;16:e01390–25. 10.1128/mbio.01390-25.

32 Hayashi S, McMahon AP. Efficient recombination in diverse tissues by a tamoxifen-inducible form of Cre: a tool for temporally regulated gene activation/inactivation in the mouse. Dev Biol 2002;244:305– 18. 10.1006/dbio.2002.0597.

33 An R, Xie E, Binns J, Rey FE, Kendziorski C, Thibeault SL. Gut-larynx axis and its contribution to laryngeal immunity. mSystems 2025;10:e01044–25. 10.1128/msystems.01044-25.

34 Borlikova G, Endo S. Inducible cAMP Early Repressor (ICER) and Brain Functions. Mol Neurobiol 2009;40:73–86. 10.1007/s12035-009-8072-1.

35 Ruchaud S, Seité P, Foulkes NS, Sassone-Corsi P, Lanotte M. The transcriptional repressor ICER and cAMP-induced programmed cell death. Oncogene 1997;15:827–36. 10.1038/sj.onc.1201248.

36 Madison BB, Dunbar L, Qiao XT, Braunstein K, Braunstein E, Gumucio DL. Cis elements of the villin gene control expression in restricted domains of the vertical (crypt) and horizontal (duodenum, cecum) axes of the intestine. J Biol Chem 2002;277:33275–83. 10.1074/jbc.M204935200.

37 Kono M, Yoshida N, Maeda K, Tsokos GC. Transcriptional factor ICER promotes glutaminolysis and the generation of Th17 cells. Proceedings of the National Academy of Sciences 2018;115:2478–83. 10.1073/pnas.1714717115.

38 Kono M, Yoshida N, Maeda K, Skinner NE, Pan W, Kyttaris VC, et al. Pyruvate dehydrogenase phosphatase catalytic subunit 2 limits Th17 differentiation. Proc Natl Acad Sci U S A 2018;115:9288–93. 10.1073/pnas.1805717115.

39 Jia L, Jiang Y, Wu L, Fu J, Du J, Luo Z, et al. Porphyromonas gingivalis aggravates colitis via a gut microbiota-linoleic acid metabolism-Th17/Treg cell balance axis. Nat Commun 2024;15:1617. 10.1038/s41467-024-45473-y.

40 Saleh MM, Frisbee AL, Leslie JL, Buonomo EL, Cowardin CA, Ma JZ, et al. Colitis-induced Th17 cells increase the risk for severe subsequent Clostridium difficile infection. Cell Host Microbe 2019;25:756-765.e5. 10.1016/j.chom.2019.03.003.

41 Pandiyan P, Bhaskaran N, Zou M, Schneider E, Jayaraman S, Huehn J. Microbiome Dependent Regulation of Tregs and Th17 Cells in Mucosa. Front Immunol 2019;10:426. 10.3389/fimmu.2019.00426.

42 Brockmann L, Tran A, Huang Y, Edwards M, Ronda C, Wang HH, et al. Intestinal microbiota-specific Th17 cells possess regulatory properties and suppress effector T cells via c-MAF and IL-10. Immunity 2023;56:2719-2735.e7. 10.1016/j.immuni.2023.11.003.

43 Burrello C, Garavaglia F, Cribiù FM, Ercoli G, Lopez G, Troisi J, et al. Therapeutic faecal microbiota transplantation controls intestinal inflammation through IL10 secretion by immune cells. Nat Commun 2018;9:5184. 10.1038/s41467-018-07359-8.

44 Busbee PB, Menzel L, Alrafas HR, Dopkins N, Becker W, Miranda K, et al. Indole-3-carbinol prevents colitis and associated microbial dysbiosis in an IL-22-dependent manner. JCI Insight 2020;5:e127551. 127551. 10.1172/jci.insight.127551.

45 Okayasu I, Hatakeyama S, Yamada M, Ohkusa T, Inagaki Y, Nakaya R. A novel method in the induction of reliable experimental acute and chronic ulcerative colitis in mice. Gastroenterology 1990;98:694–702. 10.1016/0016-5085(90)90290-H.

46 Hernández-Chirlaque C, Aranda CJ, Ocón B, Capitán-Cañadas F, Ortega-González M, Carrero JJ, et al. Germ-free and Antibiotic-treated Mice are Highly Susceptible to Epithelial Injury in DSS Colitis. J Crohns Colitis 2016;10:1324–35. 10.1093/ecco-jcc/jjw096.

47 Kong Y, Cao P, Wu J, Ye X. Prebiotics as adjunctive treatment ameliorates DSS-induced colitis and gut microbiota. Microbiology Spectrum 2025;0:e01502–25. 10.1128/spectrum.01502-25.

48 Ohl K, Nickel H, Moncrieffe H, Klemm P, Scheufen A, Föll D, et al. The transcription factor CREM drives an inflammatory phenotype of T cells in oligoarticular juvenile idiopathic arthritis. Pediatr Rheumatol Online J 2018;16:39. 10.1186/s12969-018-0253-x.

49 Zhang X, Moore CM, Harmacek LD, Domenico J, Rangaraj VR, Ideozu JE, et al. CFTR-mediated monocyte/macrophage dysfunction revealed by cystic fibrosis proband-parent comparisons. JCI Insight 2022;7:e152186. 10.1172/jci.insight.152186.

50 Li B, Alli R, Vogel P, Geiger TL. IL-10 modulates DSS-induced colitis through a macrophage – ROS – NO axis. Mucosal Immunol 2014;7:869–78. 10.1038/mi.2013.103.

51 Gu Y, Bartolomé-Casado R, Xu C, Bertocchi A, Janney A, Heuberger C, et al. Immune microniches shape intestinal Treg function. Nature 2024;628:854–62. 10.1038/s41586-024-07251-0.

52 Ohl K, Wiener A, Lippe R, Schippers A, Zorn C, Roth J, et al. CREM Alpha Enhances IL-21 Production in T Cells In Vivo and In Vitro. Front Immunol 2016;7:618. 10.3389/fimmu.2016.00618.

53 Yang B-H, Hagemann S, Mamareli P, Lauer U, Hoffmann U, Beckstette M, et al. Foxp3+ T cells expressing RORγt represent a stable regulatory T-cell effector lineage with enhanced suppressive capacity during intestinal inflammation. Mucosal Immunol 2016;9:444–57. 10.1038/mi.2015.74.

54 Hanna BS, Wang G, Galván-Peña S, Mann AO, Ramirez RN, Muñoz-Rojas AR, et al. The gut microbiota promotes distal tissue regeneration via RORγ+ regulatory T cell emissaries. Immunity 2023;56:829-846.e8. 10.1016/j.immuni.2023.01.033.

55 Zhu L, Xu L-Z, Zhao S, Shen Z-F, Shen H, Zhan L-B. Protective effect of baicalin on the regulation of Treg/Th17 balance, gut microbiota and short-chain fatty acids in rats with ulcerative colitis. Appl Microbiol Biotechnol 2020;104:5449–60. 10.1007/s00253-020-10527-w.

56 Liu H-Y, Li S, Ogamune KJ, Yuan P, Shi X, Ennab W, et al. Probiotic Lactobacillus johnsonii Reduces Intestinal Inflammation and Rebalances Splenic Treg/Th17 Responses in Dextran Sulfate Sodium-Induced Colitis. Antioxidants (Basel) 2025;14:433. 10.3390/antiox14040433.

57 Xu J, Xu H, Guo X, Zhao H, Wang J, Li J, et al. Pretreatment with an antibiotics cocktail enhances the protective effect of probiotics by regulating SCFA metabolism and Th1/Th2/Th17 cell immune responses. BMC Microbiol 2024;24:91. 10.1186/s12866-024-03251-2.

58 Britton GJ, Contijoch EJ, Mogno I, Vennaro OH, Llewellyn SR, Ng R, et al. Microbiotas from Humans with Inflammatory Bowel Disease Alter the Balance of Gut Th17 and RORγt+ Regulatory T Cells and Exacerbate Colitis in Mice. Immunity 2019;50:212-224.e4. 10.1016/j.immuni.2018.12.015.

59 Mehta P, Lawrence A, Aggarwal A. Paradoxical gastrointestinal effects of interleukin-17 blockers. Annals of the Rheumatic Diseases 2023;82:e152–e152. 10.1136/annrheumdis-2020-218719.

60 Grümme L, Dombret S, Knösel T, Skapenko A, Schulze-Koops H. Colitis induced by IL-17A-inhibitors. Clin J Gastroenterol 2024;17:263–70. 10.1007/s12328-023-01893-9.

61 Liang SC, Tan X-Y, Luxenberg DP, Karim R, Dunussi-Joannopoulos K, Collins M, et al. Interleukin (IL)-22 and IL-17 are coexpressed by Th17 cells and cooperatively enhance expression of antimicrobial peptides. J Exp Med 2006;203:2271–9. 10.1084/jem.20061308.

62 Mangan PR, Harrington LE, O’Quinn DB, Helms WS, Bullard DC, Elson CO, et al. Transforming growth factor-beta induces development of the T(H)17 lineage. Nature 2006;441:231–4. 10.1038/nature04754.

63 Aujla SJ, Chan YR, Zheng M, Fei M, Askew DJ, Pociask DA, et al. IL-22 mediates mucosal host defense against Gram-negative bacterial pneumonia. Nat Med 2008;14:275–81. 10.1038/nm1710.

64 Mitsuyama K, Niwa M, Takedatsu H, Yamasaki H, Kuwaki K, Yoshioka S, et al. Antibody markers in the diagnosis of inflammatory bowel disease. World J Gastroenterol 2016;22:1304–10. 10.3748/wjg.v22.i3.1304.

65 Ferrante M, Vermeire S, Katsanos KH, Noman M, Van Assche G, Schnitzler F, et al. Predictors of early response to infliximab in patients with ulcerative colitis. Inflamm Bowel Dis 2007;13:123–8. 10.1002/ibd.20054.

66 Esters N, Vermeire S, Joossens S, Noman M, Louis E, Belaiche J, et al. Serological markers for prediction of response to anti-tumor necrosis factor treatment in Crohn’s disease. Am J Gastroenterol 2002;97:1458–62. 10.1111/j.1572-0241.2002.05689.x.

67 Taylor KD, Plevy SE, Yang H, Landers CJ, Barry MJ, Rotter JI, et al. ANCA pattern and LTA haplotype relationship to clinical responses to anti-TNF antibody treatment in Crohn’s disease. Gastroenterology 2001;120:1347–55. 10.1053/gast.2001.23966.

68 Yoshida M, Yamada M, Sudo Y, Kojima T, Tomiyasu T, Yoshikawa N, et al. Myeloperoxidase antineutrophil cytoplasmic antibody affinity is associated with the formation of neutrophil extracellular traps in the kidney and vasculitis activity in myeloperoxidase anti-neutrophil cytoplasmic antibody-associated microscopic polyangiitis. Nephrology (Carlton) 2016;21:624–9. 10.1111/nep.12736.

69 Yoshida M, Sasaki M, Nakabayashi I, Akashi M, Tomiyasu T, Yoshikawa N, et al. Two types of myeloperoxidase-antineutrophil cytoplasmic autoantibodies with a high affinity and a low affinity in small vessel vasculitis. Clin Exp Rheumatol 2009;27:S28–32.

70 Palm NW, Zoete MR de, Cullen TW, Barry NA, Stefanowski J, Hao L, et al. Immunoglobulin A coating identifies colitogenic bacteria in inflammatory bowel disease. Cell 2014;158:1000. 10.1016/j.cell.2014.08.006.

71 Oksanen J, Simpson G, Blanchet F, Kindt R, Legendre P, Michin P, et al. _vegan: Community Ecology Package_. 2025.

