## Supplemental Information for "REGULATION OF COLONIC MACROPHAGES AND TYPE-17 AND REGULATORY T CELLS IN DSS-COLITIS BY IBD-ASSOCIATED TRANSCRIPTION FACTOR, CREM"

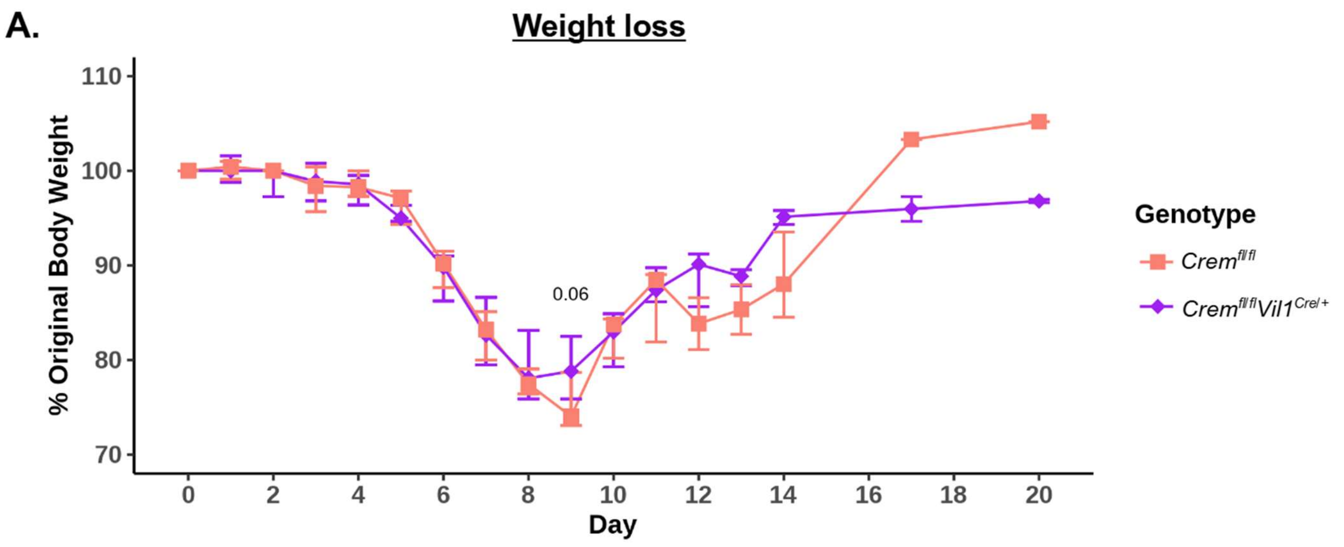


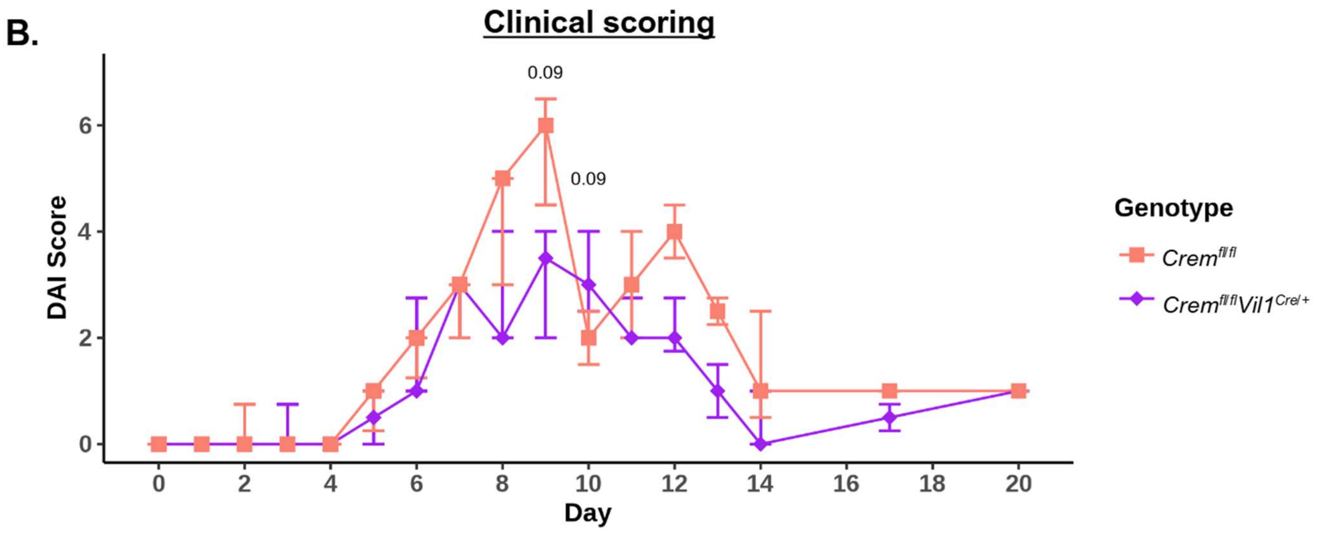


***Supplemental Figure 1. Constitutive Crem deletion in intestinal epithelial cells does not impact DSS-colitis severity.*** **A.** Percent weight loss of *Crem*$^{fl/fl}$ and *Crem*$^{fl/fl}$*Vil1*$^{Cre/+}$ mice during DSS-colitis. B. Disease activity index of *Crem*$^{fl/fl}$ and *Crem*$^{fl/fl}$*Vil1*$^{Cre/+}$ mice during DSS-colitis, scored on diarrhea, posture, and activity. **A-B.** Combined data from two independent experiments per comparison are presented as median + IQR, n = 11 per group. Statistics are calculated from a Linear Mixed Model, where % Original Body Weight or DAI is explained by an interaction between fixed effects, genotype and day, and random effect being the observation per mouse. * $p \le 0.05$, ** $p \le 0.01$.

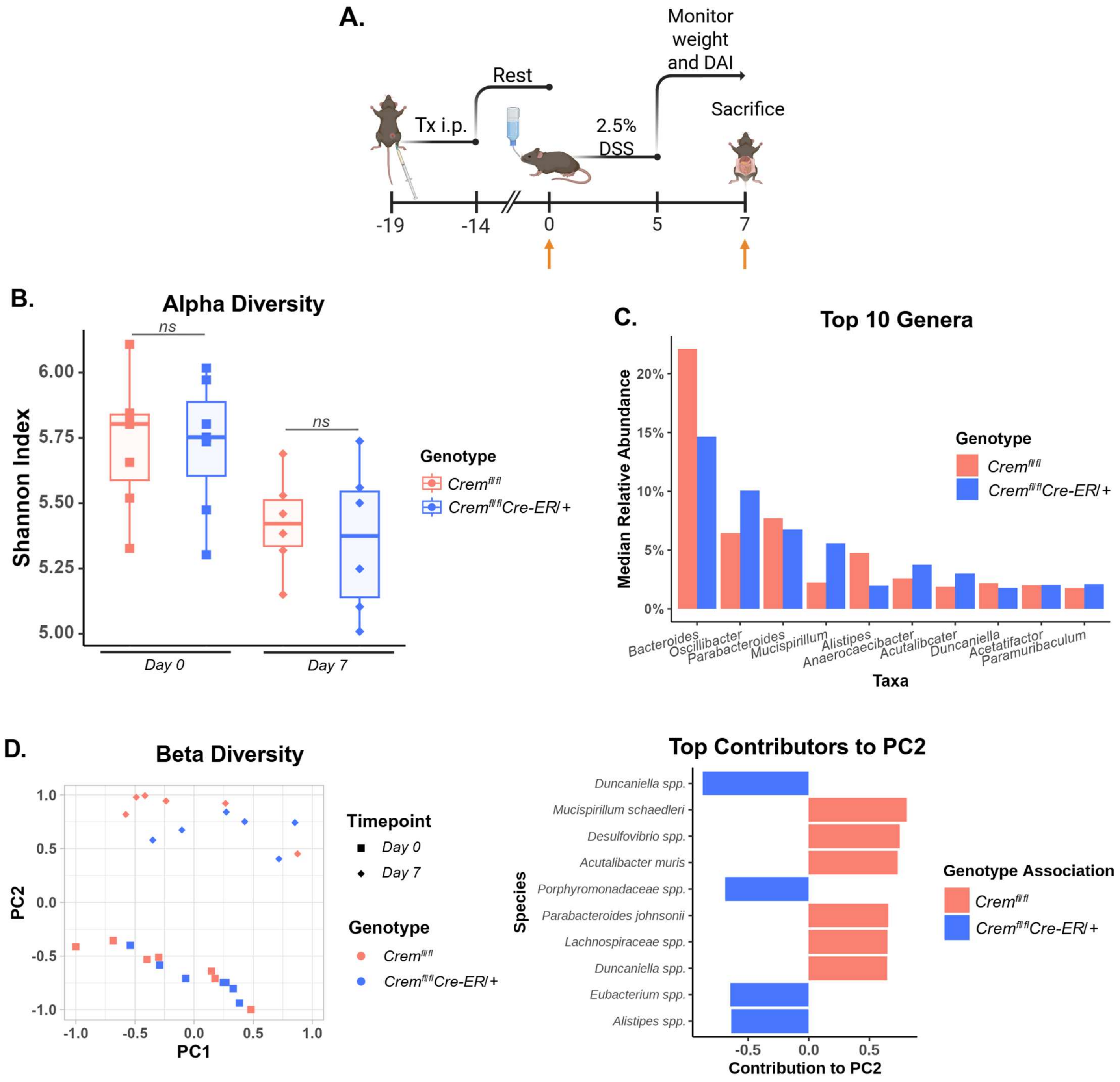


***Supplemental Figure 2.*** **Inducible global *Crem deletion* does not impact the fecal bacteriome at steady-state or during DSS-induced dysbiosis. A.** Schematic of experiment, where orange arrows indicate where stool was taken. Tx, Tamoxifen; DAI, disease activity index. **B.** Shannon index as a measure of alpha diversity**. B.** Mean relative abundance of top ten genera. **C.** Bray-Curtis dissimilarity index as a measure of beta diversity (left) and top contributors to PC2 (right). **B-D.** n = 6 per group.

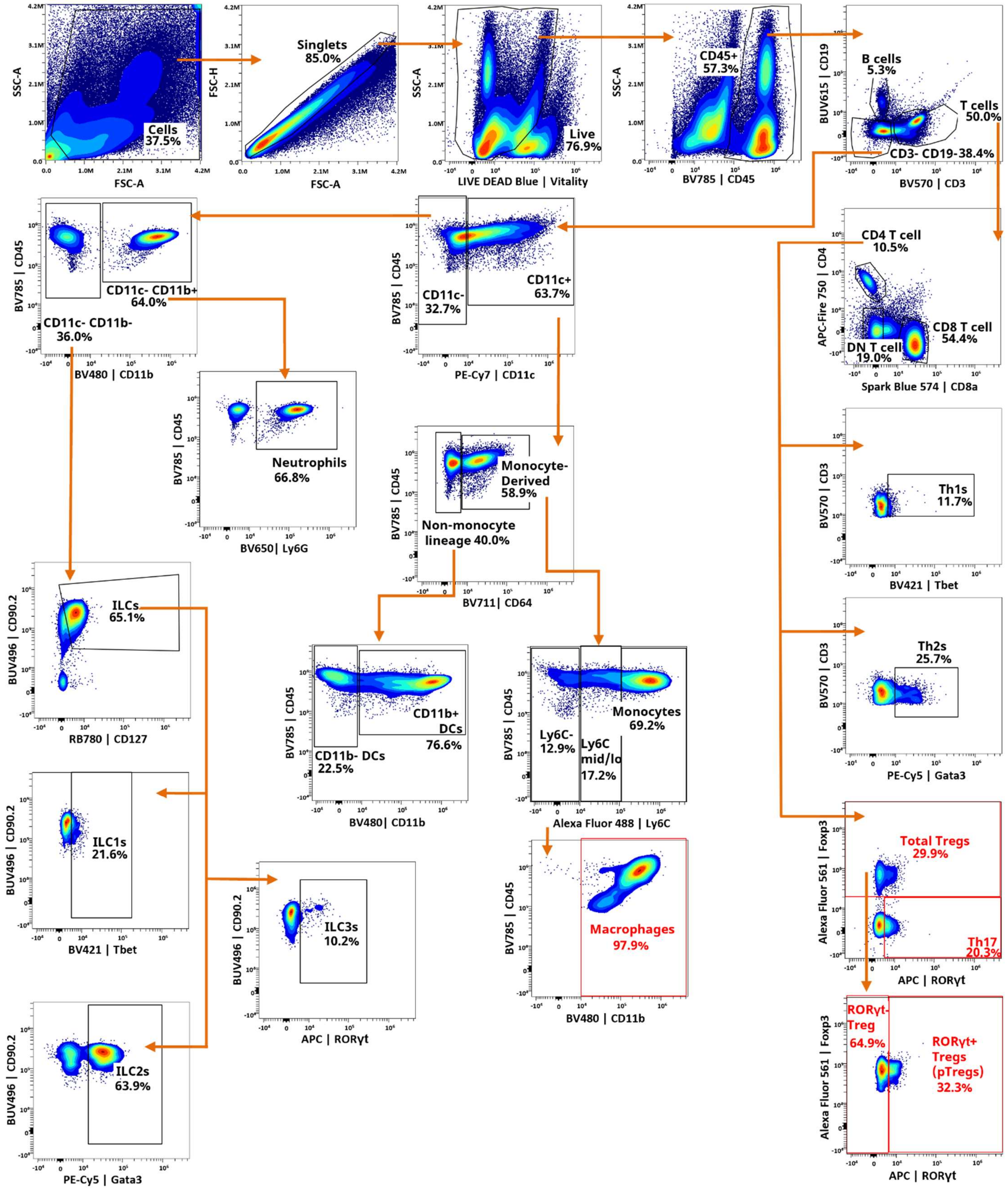


***Supplemental Figure 3. Spectral flow cytometry gating strategy for colon lamina propria.*** Populations of interest are highlighted in red.

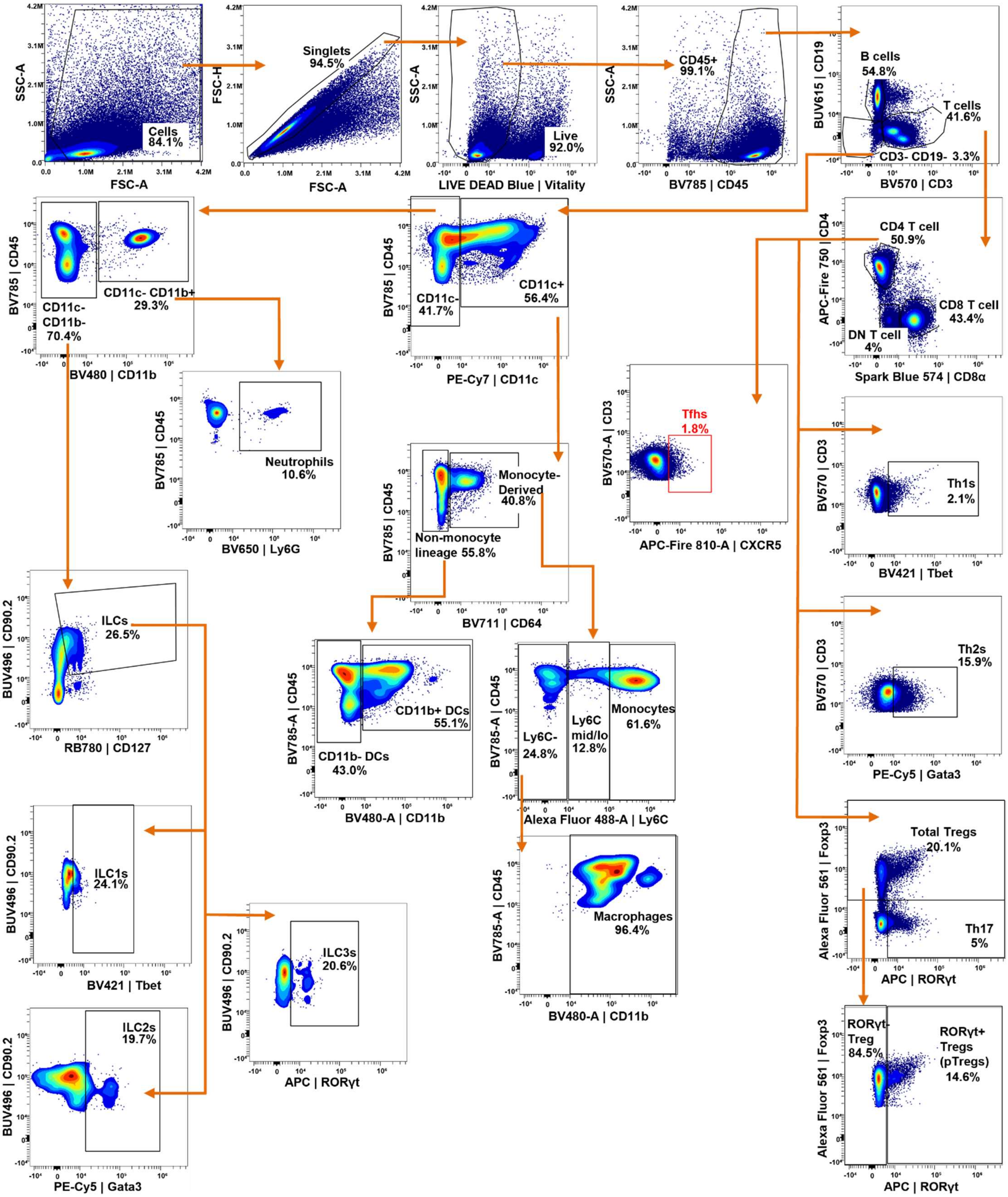


***Supplemental Figure 4. Spectral flow cytometry gating strategy for mesenteric lymph node.*** Population(s) of interest are highlighted in red where applicable.

| Marker | Fluorochrome | Dilution | Vendor/Catalog |
|---|---|---|---|
| CD3 | BV570 | 40 | BioLegend 100225 |
| CD4 | APC-Fire 750 | 100 | BioLegend 100460 |
| CD8α | Spark Blue 574 | 20 | BioLegend 100794 |
| CD11b | BV480 | 100 | BD 566117 |
| CD11c | PE-Cy7 | 33 | BioLegend 117318 |
| CD19 | BUV615 | 20 | Thermo Scientific 366-0193-82 |
| CD45 | BV785 | 40 | BioLegend 103149 |
| CD64 | BV711 | 50 | BioLegend 139311 |
| CD90.2 | BUV496 | 133 | BD 741047 |
| CD127 | RB780 | 133 | BD 569065 |
| CXCR5 | APC-Fire 810 | 80 | BioLegend 145546 |
| Ly6C | AF488 | 100 | BioLegend 128022 |
| Ly6G | BV650 | 100 | BioLegend 127641 |
| Siglec-F | AF700 | 100 | Thermo Scientific 56-1702-80 |
| Foxp3 | AF561 | 33 | Thermo Scientific 505-5773-82 |
| Gata3 | PE-Cy5 | 33 | Thermo Scientific 15996642 |
| RORγT | APC | 33 | Thermo Scientific 17-6981-82 |
| Tbet | BV421 | 25 | BD 563318 |
| Vitality | Live Dead Blue | 100 | Thermo Scientific L23105 |

***Supplemental Table 1. Comprehensive immune cell spectral flow cytometry panel.*** Table lists markers, conjugated fluorochromes, dilutions, and used to identify immune cell populations.

| ***Criteria*** | **0** | **1** | **2** | **3** | 4 |
|---:|---|---|---|---|---|
| *Weight loss* | None | < 10% | 10 – 15% | 15 – 20% | 20 – 25% |
| *Activity* | Normal | Alert / Slow-moving | Lethargic / Shaky | Inactive unless prodded | Not moving |
| *Posture* | Normal | Back-slanted | Hunched | Hunched / Nose down | - |
| *Stool consistency* | Normal | Soft stool; discolored (yellow) | Wet /stained backside / mucous | Diarrhea / No stool | - |

***Supplemental Table 2. Colitis scoring.*** Table describes scoring criteria for mice undergoing DSS-treatment and subsequent colitis.

13
